# Girls’ attentive traits associate with cerebellar to dorsal attention and default mode network connectivity

**DOI:** 10.1101/499574

**Authors:** Christiane S. Rohr, Dennis Dimond, Manuela Schuetze, Ivy Y. K. Cho, Limor Lichtenstein-Vidne, Hadas Okon-Singer, Deborah Dewey, Signe Bray

## Abstract

Attention traits are a cornerstone to the healthy development of children’s performance in the classroom, their interactions with peers, and in predicting future success and problems. The cerebellum is increasingly appreciated as a region involved in complex cognition and behavior, and moreover makes important connections to key brain networks known to support attention: the dorsal attention and default mode networks (DAN; DMN). The cerebellum has also been implicated in childhood disorders affecting attention, namely autism spectrum disorder (ASD) and attention deficit hyperactivity disorder (ADHD), suggesting that attention networks extending to the cerebellum may be important to consider in relation to attentive traits. Yet, direct investigations into the association between cerebellar FC and attentive traits are lacking. Therefore, in this study we examined attentive traits, assessed using parent reports of ADHD and ASD symptoms, in a community sample of 52 girls aged 4-7 years, i.e. around the time of school entry, and their association with cerebellar connections with the DAN and DMN. We found that cortico-cerebellar functional connectivity (FC) jointly and differentially correlated with attentive traits, through a combination of weaker and stronger FC across anterior and posterior DAN and DMN nodes. These findings suggest that cortico-cerebellar integration may play an important role in the manifestation of attentive traits.

## 1. INTRODUCTION

Children’s academic performance and interactions with peers rely heavily on attention skills (Andrade, et al., 2012; Semrud-Clikeman, 2010; Valiente, et al., 2011). Attention problems may mean that children have difficulty staying on task and engaging for longer periods of time, or have trouble screening out irrelevant stimuli, in order to focus on the information that is important (Ferretti, et al., 2008; Franceschini, et al., 2012). Poor attentional control places children at a disadvantage from the time of school entry, which can have lifelong consequences on academic attainment, employment, and social skills (Rueda, et al., 2010; Stevens and Bavelier, 2012).

Clinical manifestations of attention problems are seen in attention-deficit/hyperactivity disorder (ADHD), a neurodevelopmental disorder (NDD) with two main symptom categories: inattention and hyperactivity (Elton, et al., 2014; Larsson, et al., 2012; Matthews, et al., 2014). Attention is also frequently affected in children with autism spectrum disorder (ASD), for example, children with ASD can display extreme attention to detail and may have trouble switching attention from one task to another (Auyeung, et al., 2008). Many children with ASD are also diagnosed with ADHD (Christakou, et al., 2013; Di Martino, et al., 2013; Ray, et al., 2014) and attention problems are indeed common to several NDDs, including fetal alcohol spectrum disorder (Kooistra, et al., 2010), fragile X syndrome (Quintero, et al., 2014), Williams syndrome (Atkinson and Braddick, 2011), and Turner syndrome (Quintero, et al., 2014).

Attentive traits exist along a continuum in typically developing (TD) individuals and those with ADHD (Chabernaud, et al., 2012; Elton, et al., 2014; Elton, et al., 2016). Sampling individuals across the spectrum of these traits can therefore provide critical insight into their neural basis (Elton, et al., 2014; Forster and Lavie, 2016). Moreover, since attention problems often become apparent around the time of school entry, investigating inter-individual variability in attentive traits in a community sample of young children may elucidate the patterns that underlie these traits in childhood, and support the development of early interventions.

A growing body of literature suggests that in addition to cortico-cortico interactions (Corbetta and Shulman, 2002; Fox, et al., 2006; Rohr, et al., 2017), cortico-cerebellar connectivity may play an important role in attention (Baumann and Mattingley, 2014; Kellermann, et al., 2012; Striemer, et al., 2015). Historically, the human cerebellum has been primarily associated with sensorimotor functions (Heck and Sultan, 2002; Marr, 1969), however, evidence has accumulated regarding its role in working memory, executive functioning, language, emotion and social cognition (Schmahmann, 2010; Stoodley and Schmahmann, 2009; Strick, et al., 2009). The cerebellum is affected in NDDs affecting attention such as ADHD and ASD (Kucyi, et al., 2015; Stoodley, 2016; Wang, et al., 2014), further supporting its potential involvement in attentive traits.

The cerebellum outputs to multiple regions in the parietal and prefrontal cortices (Strick, et al., 2009), and as such, is in a unique position to influence activity in circuits implicated in attentive traits. Several functional networks involved in attention share intrinsic functional connectivity (FC) with distinct cerebellar regions, including the dorsal attention network (DAN), which is centered on the intraparietal sulcus (IPS) and the putative human frontal eye fields (FEF), and the default mode network (DMN), which is centered on the anterior cingulate cortex (ACC) and the precuneus (PCN) (Fox, et al., 2006; Habas, et al., 2009; Krienen and Buckner, 2009; O’Reilly, et al., 2010). The cortical DAN and DMN competitively interact, such that when the DAN is active the DMN is suppressed and vice versa (Buckner, et al., 2011; Corbetta and Shulman, 2002; Fox, et al., 2006). The ability to modulate or suppress DMN activity has been linked to greater top-down, goal-directed attention (Anticevic, et al., 2012; Rubia, 2013). Functionally parcellating the cerebellum is an active area of research (Wang, et al., 2016). One significant advancement was the discovery of an area in cerebellar lobule VII as a node of the DAN; this node displays the same strong negative correlation with the DMN as do cortical regions of the DAN (Brissenden, et al., 2016; Buckner, et al., 2011).

Little is known about how cortico-cerebellar FC relates to attentive traits in early childhood, when the brain undergoes vast development and remodeling. FC is often assessed using “task-free” or “resting-state” conditions wherein participants lie quietly in the MRI with no external stimulation. For young children, who naturally move more than adults, this is such a demanding task that most studies using resting-state fMRI in children past infancy begin at age 7 or 8 years (Alexander-Bloch, et al., 2013; Dosenbach, et al., 2010; Grayson and Fair, 2017; Yang, et al., 2014), with few exceptions (de Bie, et al., 2012). Part of the difficulty is that small head movements, that may be tolerable in task-based fMRI data, cause spurious signal artifacts in FC data (Power, et al., 2012; Van Dijk, et al., 2012). To improve data quality and avoid sedation during pediatric scans, labs and hospitals have been showing cartoon shows for years (Raschle, et al., 2009). It has been shown in children aged 3-11 years that head motion was significantly lower in children who were passively watching a show, than even during an age-appropriate task (Cantlon and Li, 2013; Vanderwal, et al., 2015), and several research groups have since successfully implemented and published passive viewing protocols for scanning young, awake children (Blankenship, et al., 2017; Emerson, et al., 2015; Long, et al., 2017; Rohr, et al., 2018; Rohr, et al., 2017).

In the present study, we investigated the relationship between cortico-cerebellar DAN and DMN FC, and attentive traits in a community sample of female children aged 4-7 years using a passive viewing protocol. This allowed us to assess a relatively homogenous sample that is (a) understudied with respect to the traits measured, (b) at a time when both the cerebellum (Tiemeier, et al., 2010) and cortical networks (Rohr, et al., 2017) show profound maturation, and (c) around the age of school entry when attention problems often become apparent. Given that increased intra-network FC has often been associated with greater maturity (Marek, et al., 2015; Rohr, et al., 2018), and the postulated developmental delay in brain maturation in ADHD (Shaw, et al., 2012), we hypothesized that increased difficulties in attention would be associated with weaker FC between the cerebellum and cortical regions of the DAN and DMN.

## 2. METHODS

### 2.1 Participants

Sixty-nine girls between the ages of 4 and 7 years were recruited to participate in the study. The study was approved by the Conjoint Health Research Ethics Board of the University of Calgary and performed at the Alberta Children’s Hospital. Informed consent was obtained from the parents and informed assent from the participants. Potential participants were excluded if they had a history of neurodevelopmental or psychiatric disorders, neurological problems or other medical problems that prevented participation. Participants’ data were evaluated for outliers in behavioral scores (scores > 3.5 SD from the mean) and motion on the fMRI scans (<10 minutes of usable data as described in detail below). A total of 17 participants were excluded: 3 were unable to successfully complete a practice session in an MR simulator, 9 had excessive head motion on their fMRI scan, one had insufficient cerebellar coverage, one fell asleep during fMRI acquisition, two were missing behavioral scores, and one received a diagnosis of ADHD after her visit. Therefore, the final sample consisted of 52 participants (mean age = 5.4 ± 0.8 SD years; mean IQ = 112 ±10 SD). Six of these participants were described as non right-handed by their parents: 1 was described as left-handed, 3 as more left-handed than right-handed, and 2 were described as ambidextrous. Handedness was therefore included as a regressor of no interest in the fMRI analysis.

### 2.2 Data Acquisition: Procedure

Attentive trait scores, full-scale IQ, and MR imaging data, were collected over two separate two-hour sessions within two weeks as part of a larger study. The first session consisted of several cognitive tests (reported in (Rohr, et al., 2017) including an IQ assessment using the Wechsler Preschool and Primary Scale of Intelligence - 4th Edition CDN (WPPSI-IV) (Wechsler, 2012), as well as training in an MRI simulator in order to acquaint the children with the MR environment. As showing videos increases compliance in young children (Raschle, et al., 2009), children watched an 18-minute video during training in the simulator and practiced lying still while the sounds of MR scanning were played to them via headphones. The same video (consisting of clips from the children’s TV show “Elmo’s world”) was later played for them during the real scan. If children were not comfortable in the simulator or not able to hold still, no fMRI data collection was undertaken. The actual MR scanning was undertaken in a second session.

### 2.3 Assessment of attentive traits

Attentive trait scores were obtained via validated parent reports with high internal consistency and good to acceptable test-retest and interrater reliability, respectively (4^th^ revision of the Swanson, Nolan and Pelham Parent Rating Scale (SNAP-IV, α=0.94; r=0.46): (Bussing, et al., 2008); Autism Spectrum Quotient: Children’s Version (AQ-Child, α=0.97; r=0.85): (Auyeung, et al., 2008)). Although subjective, parent reports arguably serve as a valid index of real life problems that are difficult to measure within one visit to the laboratory. The SNAP-IV includes two scales that consist of 9 items each; one scale assesses inattentive traits and the other assesses hyperactive traits. The AQ-Child includes five subscales (attention switching, attention to detail, social skills, communication and imagination). Each subscale consists of 10 items; here, we computed and utilized scores for attention switching and attention to detail. On each scale, higher scores are indicative of greater trait expression (e.g. more inattention, more difficulty switching attention).

### 2.4 Functional Connectivity

#### 2.4.1 Data Acquisition

Data were acquired on a 3T GE MR750w (Waukesha, WI) scanner using a 32-channel head coil. Functional images were acquired in 35 axial slices using a gradient-echo EPI sequence (437 volumes, TR=2500 ms, TE=30 ms, FA=70, matrix size 64×64, voxel size 3.5×3.5×3.5 mm^3^; duration: ∼18 minutes). Children watched a series of clips from “Elmo’s World” inside the MRI for the duration of the functional scan. Wakefulness was monitored using an SR-Research EyeLink 1000 (Ottawa, ON) infrared camera. Anatomical scans were acquired using a T1-weighted 3D BRAVO sequence (TR=6.764 ms, TE=2.908 ms, FA=10, matrix size 300×300, voxel size 0.8×0.8×0.8 mm^3^).

#### 2.4.2 Data Preprocessing

Data preprocessing followed a pipeline outlined by (Power, et al., 2014) and used functions from both AFNI (Cox, 1996) and FSL (Smith, et al., 2004). The pipeline included slice-time correction, motion correction and normalization to the McConnell Brain Imaging Center NIHPD asymmetrical (natural) pediatric template optimized for ages 4.5 to 8.5 years followed by normalization to 2 x 2 x 2 mm MNI152 standard space. Images were de-noised by regressing out the six motion parameters, as well as signal from white matter, cerebral spinal fluid and the global signal, and their first-order derivatives. Motion-corrupted volumes were identified by FSL MotionOutliers and censored, specifically, those that exceeded both frame-wise displacement 0.2mm and 0.3% change in BOLD signal intensity (mean censored volumes=88.6 ± 56 SD; range 2-196). It has recently been shown that this censoring approach vastly reduces motion artifacts (Power, et al., 2012; Power, et al., 2015), particularly when combined with GSR (Parkes, et al., 2018). Given that the global signal relates strongly to respiratory and other motion-induced signals, which persist through common denoising approaches (including ICA and models that attempt to approximate respiratory variance; (Power, et al., 2018), regression of the global signal was deemed necessary in this sample of young children, despite its use being controversial (Murphy, et al., 2009). Six scans exhibited a relatively large displacement (>6mm; mean 2.9mm ± 2.6 SD, range 0.6-12mm) that resulted from one large movement. As these participants contributed enough usable data (> 10min), and they were not outliers in any analysis, their data were not excluded. Head motion did not significantly correlate with age or attentive trait scores (all p>0.2). Data were spatially smoothed using a 6 mm full-width at half-maximum Gaussian kernel.

#### 2.4.3 fMRI Analysis

It was recently shown that the cerebellar node of the DAN in Crus I displays the same anti-correlation with the DMN as its cortical counterparts (Brissenden, et al., 2016). As both the DAN and DMN play important roles in attention, we reasoned that both positive and negative interactions with the cerebellum may be important. The FC analysis in this study therefore first identified the cerebellar region that showed both a positive correlation with the DAN, and a negative correlation with the DMN. This region was then used in a seed-to-voxel FC analysis, wherein FC between the cerebellar region and peaks in cortical DAN and DMN were regressed against traits of interest. To identify the cerebellar DAN node, we used two parcellation units (Yeo, et al., 2011) - one that encompassed the cortical DAN, and one that encompassed the cortical DMN (Figure 1a) - as the two seed regions of interest and computed cortical DAN and DMN FC to the cerebellum. Specifically, the average time course for each seed was extracted and entered into a voxel-wise correlation with every voxel in the cerebellum. Resultant cortico-cerebellar correlation maps were normalized using Fisher’s r-to-z transform (z=.5[ln(1+r) - ln(1-r)]) for comparison across individuals. Group-level statistical testing was conducted using a mixed-effects analysis as implemented in FSL’s FEAT. We first examined average group FC across the DAN and DMN seeds and the cerebellum using a model that contained the group intercept, z-scored handedness and the z-scored number of censored motion-affected volumes (as covariates of no interest). Voxel-wise thresholding was set at z-score > 3.1 and cluster correction was conducted using Gaussian Random Field theory with p < 0.01. A conjunction analysis between the DAN’s positive cerebellar FC map and the DMN’s negative cerebellar FC map revealed a bilateral cerebellar DAN node in Crus I (Figure 1b).

**Figure 1.**
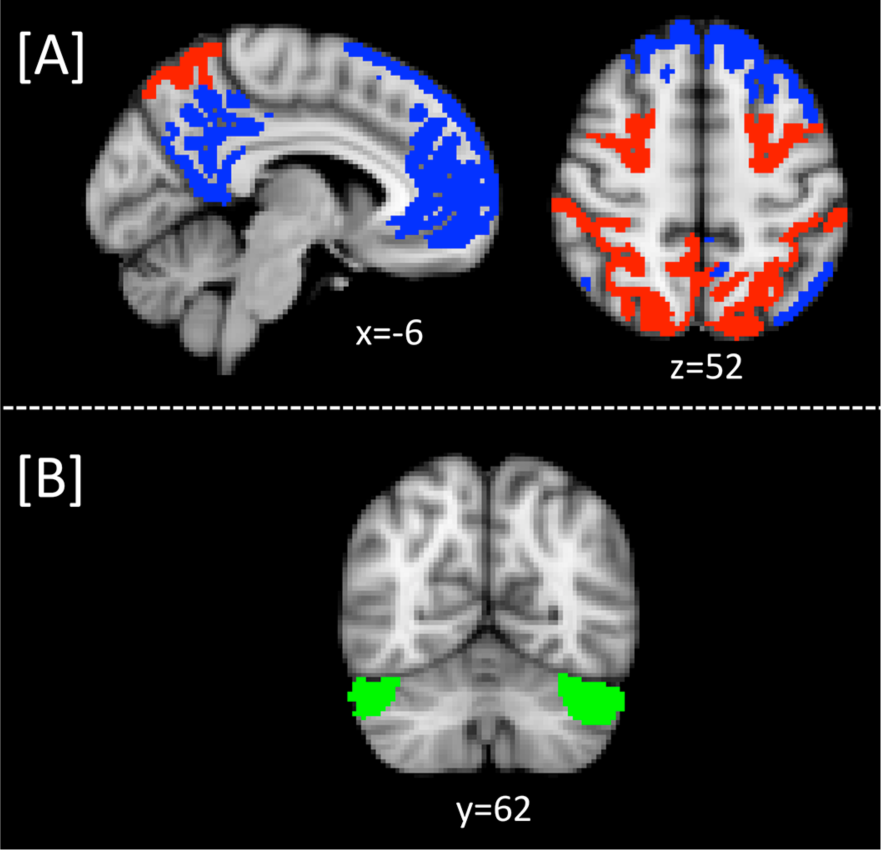
Seed regions in the cortical dorsal attention network (DAN; red) and default mode network regions (DMN; blue) displayed in panel [A] were used to identify the cerebellar DAN node by computing cortical DAN and DMN functional connectivity (FC) to the cerebellum. A conjunction analysis between the DAN’s positive cerebellar FC map and the DMN’s negative cerebellar FC map revealed a bilateral cerebellar node in Crus I, displayed in panel [B].

FC of this Crus I seed region was then computed back to the cortex in order to identify group-level FC peaks within the DAN and DMN to be used as target ROIs for subsequent analyses. FC peaks in the core nodes of the cortical DAN (bilateral IPS and FEF) and DMN (medial ACC and PCN) were determined by identifying local maxima within IPS and FEF parcellation units by (Wang, et al., 2015) and ACC and PCN parcellation units distributed in the Harvard-Oxford Atlas. We hypothesized that connections between the cerebellar DAN node Crus I and cortical DAN and DMN nodes would show a relationship with attentive traits. To identify the most significant connections underlying these traits, FC parameter estimates (i.e. strength of connectivity values) between Crus I and the cortical DAN and DMN nodes (restricted to an 8mm radius around the peaks) were entered into separate, stepwise multiple linear regression models with the four attentive trait domains we had assessed (inattention, hyperactivity, attention switching and attention to detail) in SPSS 22 (Chicago, IL); the attentive traits being dependent variables in the stepwise regression models. The stepwise criteria used was probability of F to enter ≤ 0.05, probability of F to remove ≥ 0.1, and all four multiple regression models controlled for age, head motion (as number of motion volumes detected) and full-scale IQ. In order to assess how these trait domains related to each other, we further followed up on this analysis by computing correlations between these trait domains, and by testing how well the connections identified in the four multiple regression models explained each of the four domains.

## 3. RESULTS

### 3.1 Attentive trait measures

The attentive trait scores obtained from parent reports for the children were: inattentive (SNAP-IV mean = 0.72 ± 0.44 SD; range 0-2.22; clinical range > 1.78 min-max: 0-3); hyperactive (SNAP-IV mean=0.81 ± 0.55 SD; range 0-2.56; clinical range > 1.44 min-max: 0-3); attention switching (AQ-Child mean = 12.2 ± 4 SD; range 0-15; min-max: 0-30); attention to detail (AQ-Child mean = 16.6 ± 4 SD; range 4-23; min-max: 0-30). A total of seven participants scored above clinical cut-offs (five on ADHD hyperactive traits; one on ASD traits; one on all three trait measures). We observed significant correlations between SNAP-IV inattentive and hyperactive (r=.65 and p<0.001); AQ-Child attention switching and attention to detail (r=.33 and p=0.008); inattention and attention switching (r=.26. p=0.03), and hyperactivity and attention switching (r=.24, p=0.04) (all p’s one-tailed). All other relationships were not significant (p>0.1).

### 3.2 FC of the cerebellar DAN node Crus I

Average positive and negative group FC maps of the cerebellar DAN node Crus I are shown in Figure 2 and details are provided in Table 1. As expected, we observed positive FC between Crus I and the cortical DAN, and negative FC between Crus I and the cortical DMN. The regional FC peaks within the DAN and DMN used for trait analyses are provided in Table 1 and illustrated in Figure 2.

**Table 1.** **FC of the cerebellar DAN node Crus I to cortical DAN and DMN ROIs.** Peaks shown in DAN (bilateral IPS and FEF) and DMN (medial ACC and PCN); ROIs were defined by (Wang, et al., 2015) and the Harvard-Oxford Atlas, respectively. Coordinates are given in MNI space. ACC=anterior cingulate cortex; BIL=bilateral; FEF=putative human frontal eye fields; L=left; Lat=laterality; IPS=intraparietal sulcus; M=medial; PCN=precuneus; R=right.

| Directionality | Lat | Connectivity | p-Value | Z-Max | x | y | z |
| --- | --- | --- | --- | --- | --- | --- | --- |
| positive | L | FEF | 0.00802 | 4.72 | -26 | -10 | 52 |
| positive | R | FEF | 0.00449 | 5.06 | 28 | -10 | 50 |
| positive | L | IPS | 0.000135 | 8.11 | -26 | -76 | 32 |
| positive | R | IPS | 0.000431 | 8.2 | 36 | -80 | 14 |
| negative | M | ACC | 5.96E-08 | 8.86 | -2 | 46 | 0 |
| negative | M | PCN | 1.79E-05 | 8.53 | -10 | -62 | 20 |

**Figure 2.**
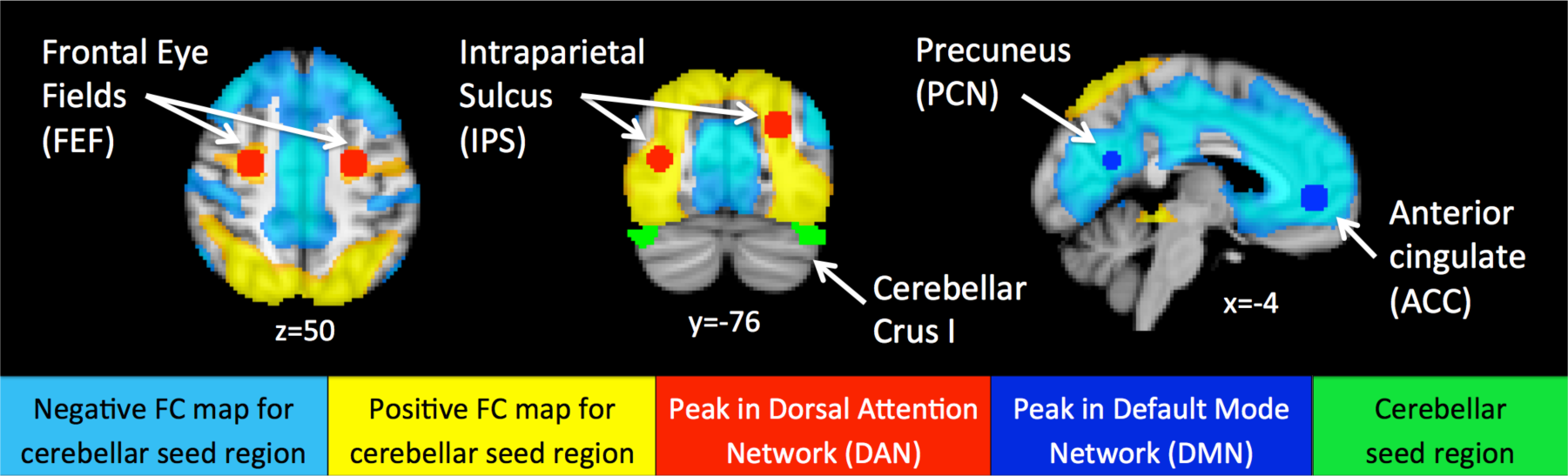
FC maps of the cerebellar DAN node Crus I, displayed in green. Positive FC is displayed in yellow and was observed between Crus I and the cortical DAN. Negative FC is displayed in light blue and was observed between Crus I and the cortical DMN. IPS and FEF peaks in the DAN are denoted in red, ACC and PCN peaks in the DMN in dark blue.

### 3.3 FC associated with attentive traits

FC between the Crus I seed and cortical DAN and DMN nodes significantly correlated with inattention and attention switching, but not hyperactivity or attention to detail. The two respective final models (significant at p=0.001 and p=0.001), which controlled for age, head motion, and IQ, identified the connections underlying these traits as follows: inattentive traits were positively associated with Crus I – ACC FC (partial r=0.35, p=0.01), meaning that the typical pattern of negative FC between Crus I and the cortical DMN node was *weaker* when inattention was more pronounced in children. Inattentive traits were also positively associated with Crus I – FEF FC (partial r =0.53, p=0.00005), meaning that positive FC between Crus I and the cortical DAN node was *stronger* when inattention was more pronounced (Figure 3). Note that partial r’s derive from the regression, and are influenced by other connections in the model (for simple partial and bivariate correlations between attentive traits and connections, see Supplementary Tables and Figures S1 and S2).

**Figure 3.**
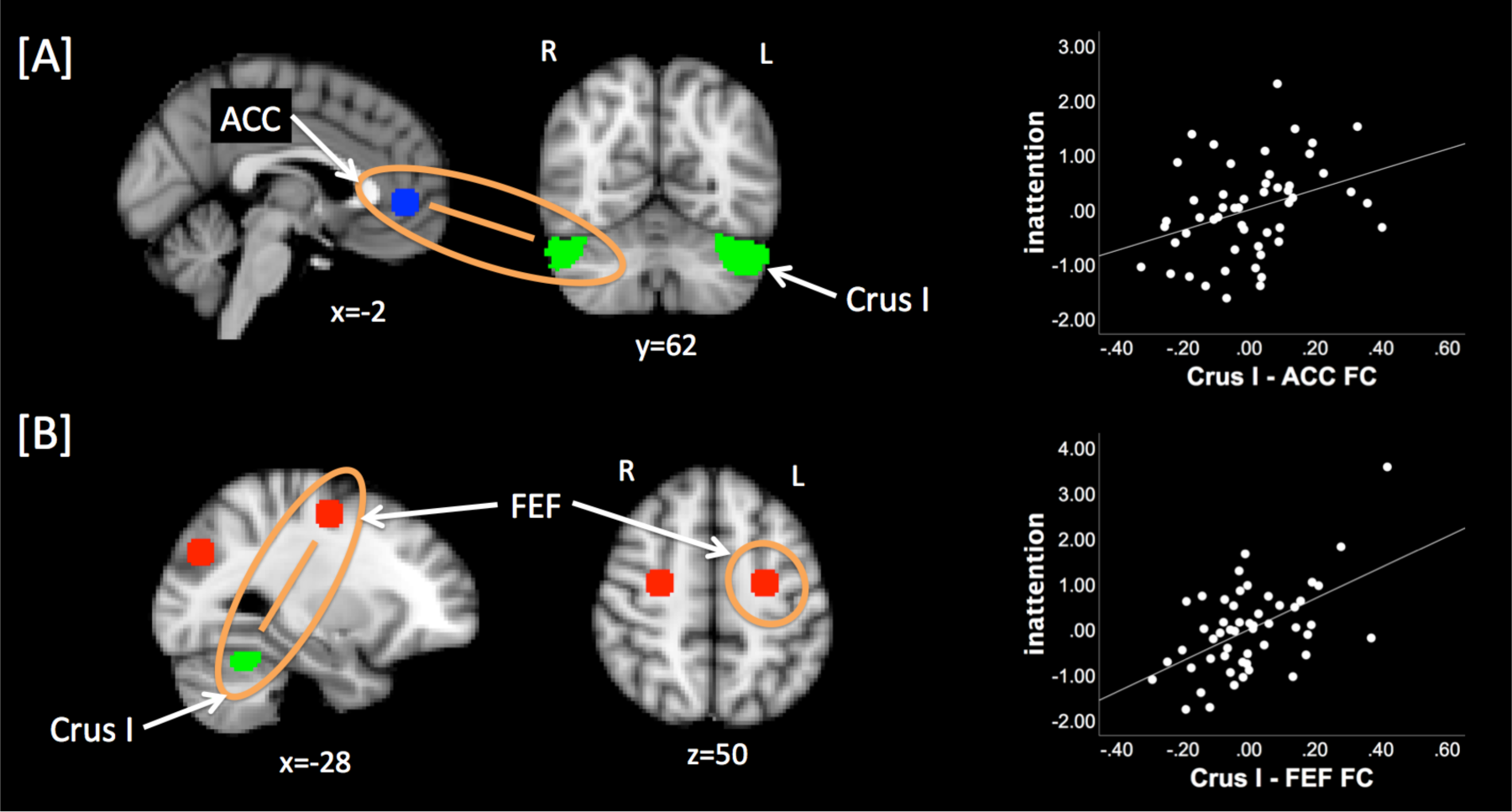
FC between the cerebellar DAN node and cortical DAN and DMN nodes correlated with inattentive traits. [A] Inattentive traits were positively associated with Crus I – ACC FC, meaning that FC between the cerebellar DAN node and the cortical DMN node was weaker when inattentive traits were more pronounced in children. [B] Inattentive traits were also positively associated with Crus I – FEF FC, meaning that FC between the cerebellar DAN node and the cortical DAN node was stronger when inattentive traits were more pronounced in children. Partial regression plots for [A] and [B] relationships are shown on the right. ACC=anterior cingulate cortex; FEF=putative human frontal eye fields; L=left; R=right.

Attention switching traits were partially reflected in an FC pattern almost identical to that for inattentive traits: we found a positive association with Crus I – ACC FC (partial r =0.37, p=0.007), as well as with Crus I – FEF FC (partial r =0.33, p=0.02), the only difference being that cerebellar FC with the right FEF node was associated with attention switching, rather than cerebellar FC with the left FEF node, which was associated with inattention. In addition, attention switching traits were negatively associated with Crus I – PCN FC (partial r=-0.34, p=0.01), meaning that negative FC between Crus I and the cortical DMN node was *stronger* when difficulties switching attention were more pronounced in children. Attention switching traits were also negatively associated with Crus I – IPS FC meaning that positive FC between Crus I and the cortical DAN node was *weaker* when difficulties switching attention were more pronounced (partial r=-0.33, p=0.02) (Figure 4).

**Figure 4.**
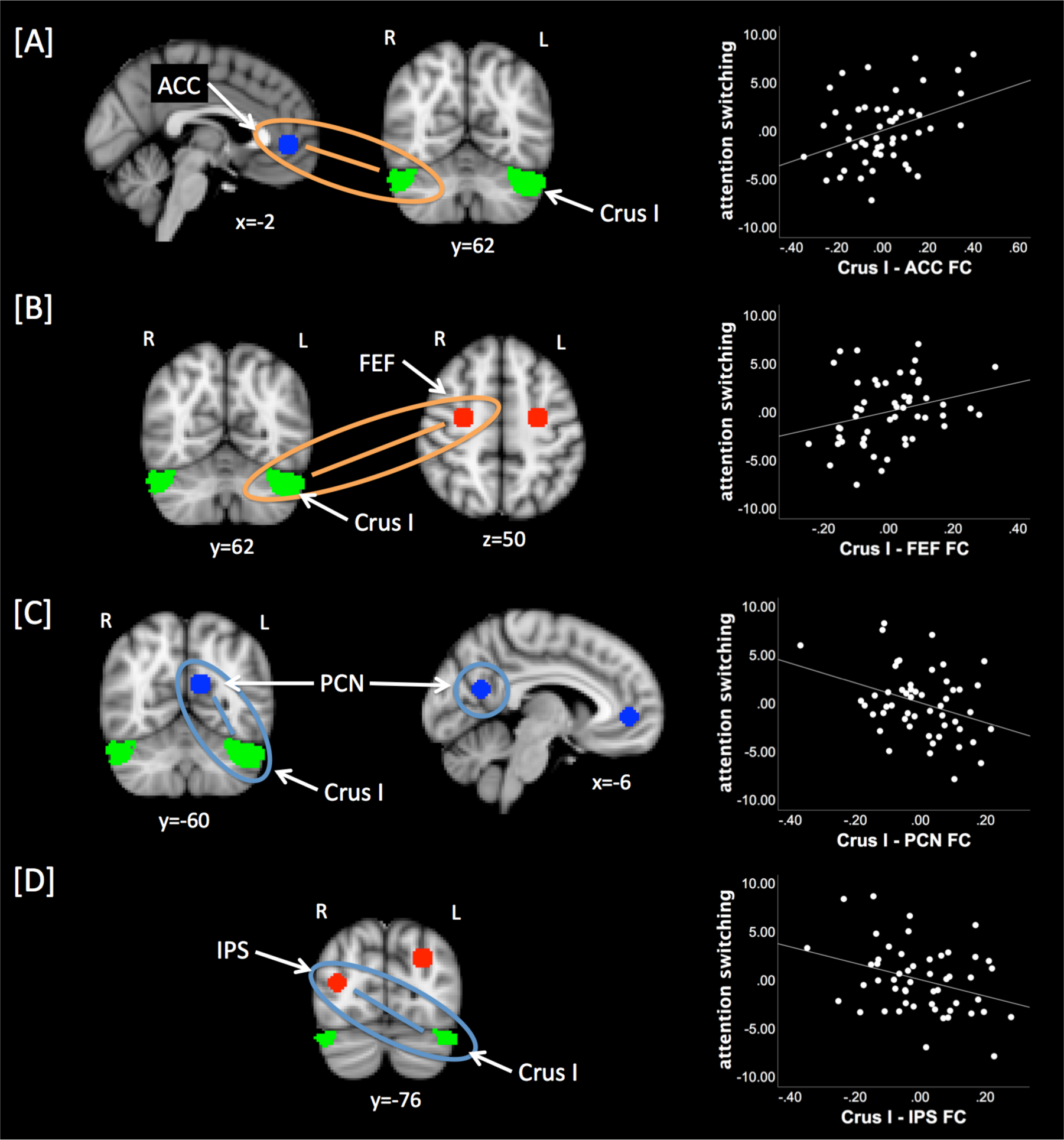
FC between crus I and cortical DAN and DMN nodes correlated with attention switching traits. Panels [A] and [B] depict FC patterns associated with attention switching that are similar to those underlying inattentive traits, with the exception that [B] involves the right FEF node. [C] Attention switching traits were negatively associated with Crus I – PCN FC, meaning that FC between crus I and the cortical DMN node was stronger when difficulties switching attention were more pronounced in children. [D] Attention switching traits were also negatively associated with Crus I – IPS FC, meaning that FC between Crus I and the cortical DAN node was weaker when difficulties switching attention were more pronounced in children. Partial regression plots for all relationships are shown on the right. L=left; IPS=intraparietal sulcus; PCN=precuneus; R=right.

A follow-up analysis testing how well the connections identified in the two multiple regression models explained each of the four attention traits revealed that the models for inattentive and attention switching traits cross-correlated (p=0.028 and p=0.01, respectively), while the remaining cross-correlations were not significant (all p>.17).

## 4. DISCUSSION

This study tested the hypothesis that patterns of cortico-cerebellar FC, in networks associated with attention, would be associated with attentive traits in early childhood. We showed that in a community sample of girls aged 4-7 years, differential patterns of FC between the cerebellar DAN node (Crus I) and cortical DAN and DMN nodes significantly correlated with inattentive traits, as well as traits related to the ability to switch attention, but not hyperactive traits or the ability to pay attention to detail. Specifically, both inattention and attention switching were positively associated with FC between Crus I and prefrontal DAN and DMN nodes. In addition, we observed that attention switching was negatively associated with FC between Crus I and parietal DAN and DMN nodes. As such, these results partially confirmed our hypothesis that weaker cortico-cerebellar FC would be associated with greater attentive problems, however, this pattern was limited. Our finding of increased cortico-cerebellar FC in children with greater attentive problems was more surprising. Together, these findings suggest a complex reality in that it might not only be weaker FC, but the *combination* of weaker and stronger cortical DAN and DMN FC with Crus I that relates to the spectrum of attentive traits in TD children. Our results are particularly interesting given that this is a dynamic period of brain development in which anterior and posterior brain regions are undergoing distinct developmental trajectories (Krongold, et al., 2017), and that both the DMN and DAN include prefrontal and parietal nodes.

Our finding that inattentive traits are associated with positive integration of Crus I with DAN and DMN, while attention switching traits are associated with a mix of positive and negative integration of Crus I with DAN and DMN, sheds new light on the neural basis of these behaviors. Stronger FC between the DAN and the DMN, which was related to weaker attention switching traits in our data, has previously been linked to more difficulties in boys with ASD in a study by Elton and colleagues (2016); notably, attention switching is one of the two attention subdomains on the ASD assessment we employed here. As this effect was not present in a categorical analysis contrasting TD boys and boys with ASD (Elton, et al., 2016), this suggests that an alteration in these networks relates to attentive difficulties that are not unique to ASD, but rather exist across the general population to varying degrees. Our work adds to these findings by showing that this relationship is not unique to boys aged 6.5-18 years, but also exists in girls as young as 4-7 years of age when probing the cortico-cerebellar DAN’s FC with the DMN.

We also found that greater FC between Crus I and a cortical DAN region was associated with greater inattentive traits. Greater activity in the DAN in relation to ADHD (of which inattention is one of the two main symptoms) has previously been shown and interpreted as compensatory effort (Cortese, et al., 2012). While several studies have reported increased DMN activity during tasks in ADHD vs. TD participants, (Sidlauskaite, et al., 2016) found that the process of down-regulating the DMN when preparing to switch from rest to task was unimpaired in ADHD. This is a somewhat counter-intuitive, as the ability to modulate or suppress DMN activity via the DAN is linked to greater top-down, goal-directed attention (Anticevic, et al., 2012; Rubia, 2013), which is an area of difficulty in ADHD. Instead, Sidlauskaite et al., (2016) found difficulties up-regulating the DMN when switching from task to rest in individuals with ADHD. It has been suggested that there may be two attentional states (Esterman, et al., 2014): one that relies on the DMN and is relatively effortless and ideal for sustaining attention (such as when resting or watching a movie), and one that relies on the DAN, is effortful and more suited to selective or executive attention (such as when performing a task). Thus, attention would alternately rely on the DAN and DMN, when operating in effortful or relatively effortless modes, respectively; in both cases, however, the DAN and the DMN would be anti-correlated. Selectively increased or decreased FC within and between the DAN and DMN in covariance with attentive traits may therefore reflect an imbalance in neural processes mediated by the cerebellum and related to attentive processing.

Our results further highlight the importance of cerebellar network FC in relation to attention problems, and are in line with recent works that have shown that attention relies on a multitude of regions not traditionally ascribed to the canonical attention networks (Rohr, et al., 2018; Rosenberg, et al., 2016). Indeed, in recent years there has been a surge in studies showcasing the cerebellum’s role in attention (Baumann and Mattingley, 2014; Kellermann, et al., 2012; Striemer, et al., 2015). Genetic, animal model, post-mortem and neuroimaging studies, have all reported cerebellar differences between individuals with ADHD and those who are TD (Arime, et al., 2011; Baroni and Castellanos, 2015; Bonvicini, et al., 2016; Chess and Green, 2008; Rubia, et al., 2014; van der Meer, et al., 2016), as well as between TD individuals and those with ASD (for reviews see (Becker and Stoodley, 2013; Crippa, et al., 2016; Stoodley, 2016; Wang, et al., 2014). To our knowledge, however, this is only the second study to describe cerebellar FC with cortical networks in relation to attention or attention problems (see (Kucyi, et al., 2015) for an investigation into cerebellar FC with the DMN in relation to ADHD).

Though few studies have assessed attention-related trait dimensions across both TD children and children with NDDs, Elton and colleagues (2014, 2016) have demonstrated in two large studies utilizing data from the Autism Brain Imaging Data Exchange and ADHD-200 Sample databases that both categorical effects of NDDs (i.e. effects specific to the NDD groups) as well as dimensional effects (traits spanning across NDDs and TD participants) exist. This highlights the promise of looking into dimensional effects of attentive traits across the TD population. Recent studies across various neuropsychological domains show that taking individual differences into account can help elucidate the neural underpinnings of the diversity and complexity of cognitive, emotional, and social traits in both neurotypical (Fuentes-Claramonte, et al., 2016; Goldfarb, et al., 2016; Rohr, et al., 2015; Rohr, et al., 2013; Rohr, et al., 2016; Vossel, et al., 2016) and clinical groups (Nebel, et al., 2015; van Dongen, et al., 2015; von Rhein, et al., 2015). The use of dimensional approaches can also allow for more statistical power in studies on ADHD and ASD, which are chronically underpowered due to heterogeneity in the populations studied (Fair, et al., 2012; Nigg, 2005; Sonuga-Barke, 2002; Sonuga-Barke, et al., 2008) and the challenge of recruiting and successfully collecting neuroimaging data from these groups.

Strengths of the study presented here include the use of four measures of attention traits in a relatively large group of young female children, at an age when attention problems typically first emerge, and a sufficiently long scan time to obtain reliable FC estimates. Although parent reports are subjective, they capture a measure of behavior integrated over a longer time frame than can be observed in a laboratory visit. There are also limitations. Current research implies sex differences in attentive traits at the extreme ends of the spectrum in individuals with ADHD and ASD (Gaub and Carlson, 1997; Rucklidge, 2008), yet assessments of attention skills in TD children indicate that such sex differences might not be present across other parts of the spectrum (Breckenridge, et al., 2013). Given that the assessments we utilized were developed based on boys, and the parallels with previous research in boys that we observe here, a generalization of our results would be plausible. However, future studies will need to test whether our findings also apply to male neural expressions of attentive traits. At the same time, some expressions specific to females, e.g. the female expression of hyperactivity, may not have been adequately captured by the SNAP-IV and the Child-AQ, which is a potential explanation for why we observed no association between Crus I FC and hyperactivity or attention to detail in the data presented here. Secondly, we regressed out the global signal to combat effects of motion in our young children’s data as per current recommendations (Parkes, et al., 2018; Power, et al., 2018) and because of doubts about the accuracy of older studies, which may not have adequately controlled for head motion (Grayson and Fair, 2017). Global signal regression (GSR) changes the distribution of FC estimates so that it is approximately centered on zero, which enhances negative correlations (Murphy, et al., 2009); there is currently no way of knowing these are generally artefactual or real, as the “true brain state” is not known (Murphy and Fox, 2017). Given that the negative correlations that we observed here between the DAN and the DMN are well established and indeed expected, they are unlikely to have been artificially introduced, but they could have been enhanced. The extent to which GSR removes noise or signal may further be contingent on the amount of global noise present (Chen, et al., 2012), meaning individual differences in head motion may cause differences in noise levels and thus lead to GSR exerting a differential effect on FC (Parkes, et al., 2018). We have therefore controlled for head motion both in the calculation of FC, as well as in the multiple regression models. Despite these efforts, potential challenges in the interpretation of our results remain. Finally, children in this study were engaged in passive viewing of video clips during functional data acquisition, rather than at rest. It has been previously shown that networks are largely similar, though not identical, during passive viewing and rest (Bray, et al., 2015; Emerson, et al., 2015; Vanderwal, et al., 2015). For instance, the spatial extent of the DAN and the DMN appears to differ (Bray, et al., 2015), as well as inter-network FC between the visual system and parts of the DAN (Emerson, et al., 2015) and the DMN (Vanderwal, et al., 2015). While we would expect this to mainly impact FC across the group rather than individual differences, and while passive viewing paradigms are an essential tool in the acquisition of usable fMRI data in children younger than 7 years of age, this may limit comparability across studies.

In summary, our findings suggest that young girls’ attentive traits are underlined by both enhanced and reduced FC patterns within the DAN, and enhanced and reduced FC between the DAN’s cerebellar node Crus I and the DMN. Our findings concur with, and tie together, a body of research that highlights the involvement of the cerebellum, as well as the DAN and DMN, in attentive traits. To our knowledge, this is the first study to investigate FC of the cerebellar DAN node in relation to attentive traits in early childhood. The results highlight insights into how the brain’s cortico-cerebellar networks are associated with children’s traits and may serve as a basis for future studies investigating a variety of attentive traits and cortico-cerebellar networks across TD and NDD populations.

## Acknowledgements

This work was supported by Natural Sciences and Engineering Research Council of Canada (NSERC); Canadian Institutes of Health Research - Institute of Neurosciences, Mental Health and Addiction (CIHR-INMHA); and Alberta Children’s Hospital Research Institute (ACHRI) grants awarded to Dr. Bray, as well as University of Calgary Eyes High, Alberta Innovates and NSERC CREATE I3T Postdoctoral Fellowships awarded to Dr. Rohr. The funders had no role in study design, data collection and analysis, decision to publish or preparation of the manuscript.

