## Supplements for "Girls’ attentive traits associate with cerebellar to dorsal attention and default mode network connectivity"

| | | l Crus - l IPS | l Crus - r IPS | r Crus - l IPS | r Crus - r IPS | l Crus - l FEF | l Crus - r FEF | r Crus - r FEF | l Crus - ACC | r Crus - ACC | l Crus - PCN | r Crus - PCN |
|---|---|---|---|---|---|---|---|---|---|---|---|---|
| Inattention | r | -0.06 | -0.09 | -0.16 | 0.08 | 0.48 | 0.40 | -0.14 | -0.36 | 0.25 | -0.25 | 0.16 |
| | p | 0.69 | 0.55 | 0.28 | 0.57 | 0.00 | 0.00 | 0.33 | 0.01 | 0.08 | 0.08 | 0.26 |
| Hyperactivity | r | -0.24 | -0.05 | -0.01 | 0.13 | 0.32 | 0.21 | 0.03 | -0.30 | 0.13 | -0.29 | -0.07 |
| | p | 0.10 | 0.74 | 0.93 | 0.37 | 0.02 | 0.15 | 0.86 | 0.04 | 0.39 | 0.05 | 0.64 |
| Attention Switching | r | -0.23 | -0.31 | -0.14 | -0.19 | 0.18 | 0.26 | -0.10 | -0.04 | 0.33 | -0.31 | 0.06 |
| | p | 0.12 | 0.03 | 0.34 | 0.20 | 0.22 | 0.08 | 0.50 | 0.78 | 0.02 | 0.03 | 0.67 |
| Attention to Detail | r | -0.31 | -0.22 | 0.26 | 0.18 | 0.11 | -0.04 | -0.03 | -0.06 | 0.12 | -0.24 | 0.25 |
| | p | 0.03 | 0.13 | 0.08 | 0.22 | 0.44 | 0.81 | 0.82 | 0.68 | 0.42 | 0.10 | 0.09 |

**Supplementary Table S1. Partial correlations between attentive traits and cortico-cerebellar FC nodes within DAN and DMN, controlling for age, IQ and motion.** P-values are uncorrected. ACC=anterior cingulate cortex; FEF=putative human frontal eye fields; l=left; IPS=intraparietal sulcus; PCN=precuneus; r=right.

| | | l Crus - l IPS | l Crus - r IPS | r Crus - l IPS | r Crus - r IPS | l Crus - l FEF | l Crus - r FEF | r Crus - r FEF | l Crus - ACC | r Crus - ACC | l Crus - PCN | r Crus - PCN |
|---|---|---|---|---|---|---|---|---|---|---|---|---|
| Inattention | r | -0.04 | -0.08 | -0.16 | 0.10 | 0.44 | 0.39 | -0.17 | -0.31 | 0.25 | -0.26 | 0.14 |
| | p | 0.77 | 0.59 | 0.26 | 0.47 | 0.00 | 0.01 | 0.24 | 0.03 | 0.07 | 0.07 | 0.31 |
| Hyperactivity | r | -0.21 | -0.02 | -0.05 | 0.14 | 0.25 | 0.21 | 0.00 | -0.22 | 0.18 | -0.23 | -0.09 |
| | p | 0.15 | 0.89 | 0.72 | 0.32 | 0.08 | 0.14 | 0.99 | 0.12 | 0.20 | 0.10 | 0.51 |
| Attention Switching | r | -0.11 | -0.26 | -0.11 | -0.15 | 0.22 | 0.33 | -0.09 | -0.12 | 0.28 | -0.32 | 0.02 |
| | p | 0.45 | 0.07 | 0.43 | 0.28 | 0.13 | 0.02 | 0.52 | 0.40 | 0.05 | 0.02 | 0.88 |
| Attention to Detail | r | -0.20 | -0.17 | 0.23 | 0.19 | 0.11 | 0.05 | -0.03 | -0.08 | 0.14 | -0.21 | 0.19 |
| | p | 0.16 | 0.24 | 0.10 | 0.18 | 0.45 | 0.74 | 0.81 | 0.57 | 0.33 | 0.14 | 0.17 |

**Supplementary Table S2. Bivariate correlations between attentive traits and cortico-cerebellar FC nodes within DAN and DMN.** P-values are uncorrected. ACC=anterior cingulate cortex; FEF=putative human frontal eye fields; l=left; IPS=intraparietal sulcus; PCN=precuneus; r=right.

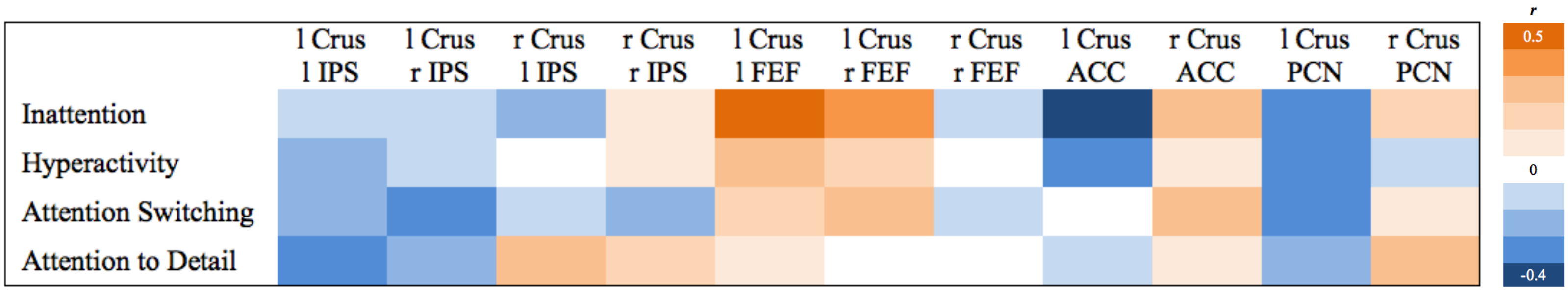


**Supplementary Figure S1. Visualisation of r-values of partial correlations between attentive traits and cortico-cerebellar FC nodes within DAN and DMN, controlling for age, IQ and motion.** ACC=anterior cingulate cortex; FEF=putative human frontal eye fields; l=left; IPS=intraparietal sulcus; PCN=precuneus; r=right.

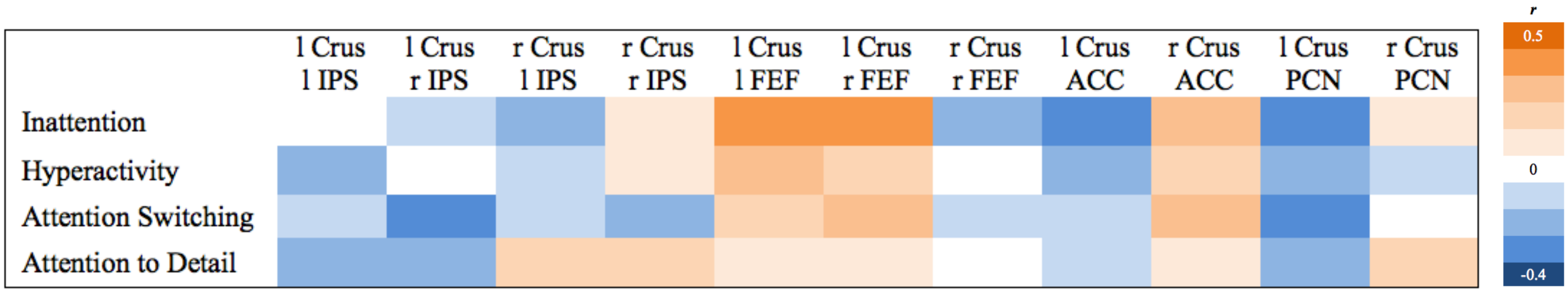


**Supplementary Figure S2. Visualisation of r-values of bivariate correlations between attentive traits and cortico-cerebellar FC nodes within DAN and DMN.** ACC=anterior cingulate cortex; FEF=putative human frontal eye fields; l=left; IPS=intraparietal sulcus; PCN=precuneus; r=right.
